# Oligomeric coiled-coil adhesins that drive chain-like adhesion diversify surface colonization strategies in Shiga toxin-producing *Escherichia coli*

**DOI:** 10.64898/2025.12.25.695649

**Authors:** Yuto Kotaka, Naoki A Uemura, Tadayuki Iwase, Ken-ichi Lee, Nozomi Ishijima, Daisuke Nakane, Hirotaka Kobayashi, Michiyo Kataoka, Yukihiro Akeda, Sunao Iyoda

## Abstract

Bacteria frequently colonize host and environmental surfaces under fluid flow. Chain-like adherence pattern (CLAP) is an EibG-mediated surface colonization phenotype of certain Shiga toxin-producing *Escherichia coli* (STEC) that lack the locus of enterocyte effacement (LEE). EibG, an immunoglobulin-binding oligomeric coiled-coil adhesin, drives CLAP, but the temporal dynamics and genetic diversity underlying chain formation remain unclear. Here, we use live-cell time-lapse imaging to show that chains arise from single cells that elongate and divide without separation. Under flow, chains resist detachment and undergo shear-dependent fragmentation at cell-cell junctions, releasing viable clonal units that disperse downstream. Comparative genomics reveals substantial diversity among EibG-related adhesins and identifies distinct lineages, including chain-like adhesins (*cla*) that mediate CLAP while lacking IgG binding. Screening of 1,354 LEE-negative STEC genomes from England shows that *claB* is present in 95.6% of strains from major LEE-negative STEC serotypes, highlighting its epidemiological prevalence. Targeted mutagenesis demonstrates that chain formation and IgG binding are mediated by distinct structural domains, revealing the modular functional architecture of these adhesins. In a mouse infection model, deletion of *eibG* reduced lethality, indicating that EibG contributes to virulence. Collectively, these findings establish CLAP as a dynamic, surface-associated strategy of LEE-negative STEC and reveal previously unrecognized diversification among adhesins that drive this behavior.

## Introduction

Shiga toxin-producing *Escherichia coli* (STEC) are major foodborne pathogens that cause severe disease outbreaks worldwide. Young children and older adults are at increased risk of developing severe diseases ^1^. Clinical manifestations range from abdominal pain and bloody diarrhea to life-threatening complications such as hemolytic uremic syndrome (HUS) and encephalopathy^2,3^. Most clinical STEC isolates harbor the locus of enterocyte effacement (LEE), which encodes the adhesin Intimin, a type III protein secretion system (T3SS), and multiple effector proteins necessary for virulence^4,5^.

Nevertheless, several large-scale outbreaks have been caused by LEE-negative STEC strains^6–8^. A notable example is the 2011 outbreak in Germany involving the LEE-negative STEC O104:H4 strain, which resulted in an unusually high rate of HUS and substantial mortality^7^. Genomic analyses later revealed that this strain represents a hybrid strain, in which an enteroaggregative *E. coli* (EAEC) lineage acquired a Shiga toxin-encoding prophage^9^. Multiple STEC virulence factor —including Shiga toxin, hemolysin, and several adhesins —are encoded on mobile genetic elements (MGEs), such as bacteriophages and plasmids, enabling horizontal transfer among diarrheagenic *E. coli* (DEC)^10,11^. Consequently, the horizontal acquisition of mobile genetic elements can convert strongly adherent DEC lineages into highly virulent pathogens, as illustrated by the emergence of the O104:H4 strain^12,13^.

In Japan, O91 is one of the most frequently isolated LEE-negative STEC serogroup from clinical cases. Similar trend has been reported in the United Kingdom and across the European Union^14,15^. Many O91:H14 isolates exhibit a distinctive chain-like adherence pattern (CLAP), mediated by EibG, an immunoglobulin-binding protein structurally related to YadA in *Yersinia enterocolitica*^16,17^. This adherence pattern is morphologically similar to the adherence pattern characteristic of EAEC, conferring strong attachment to host epithelial cells^17^. Notably, some strains forming CLAP (excluding O91:H14) possess canonical EAEC virulence markers, suggesting evolutionary and functional convergence with the EAEC lineage^18,19^.

EibG is a 508-amino acid oligomeric coiled-coil adhesin (OCA) that mediates immunoglobulin binding and CLAP^17^. Structural studies of the related protein EibD have revealed a lollipop-like structure comprising a globular head domain and right- and/or left-handed coiled-coil stalk regions^20^. EibG binds immunoglobulins in a non–immune manner by interacting with the Fc fragment of IgG^17^. Its expression is highly condition-dependent: robust expression is observed during static culture, whereas only minimal expression levels are detected under shaking conditions^21^. EibG-expressing cells also exhibit pronounced autoaggregation and biofilm formation^22^. Recent domain deletion and chimeric analyses have demonstrated that EibG-mediated chain formation requires an intact head domain as deletion of the entire head domain —rather than the N-terminal domain (NTD) or Left-handed parallel β-roll (LPBR) alone —abolishes the chain-formation phenotype^23^. However, whether chain formation and IgG Fc-binding activity are functionally linked or mediated by distinct structural domains of EibG remains unclear.

Despite the clear importance of EibG-mediated chain formation in adherence and colonization, its contribution to STEC virulence is not fully understood. In this study, we performed comprehensive analyses combining time-lapse microscopy, comparative genomics of clinical isolates, and mutant characterization to elucidate the biological functions of EibG. We further describe Cla, a newly identified CLAP factor lacking IgG Fc-binding activity, and characterized its epidemiological relevance by screening publicly available genomes from a UK surveillance cohort. These findings reveal that EibG-mediated chain formation represents a previously unrecognized surface colonization strategy in STEC and provide insights into adhesive traits that may contribute to the pathogenic potential of hybrid DEC lineages.

## Results

### A single multicellular chain originates from a single cell

STEC O91:H14 strains (including O91:H–/Hg14), displayed a characteristic CLAP when infecting HEp-2 cells. To resolve the ultrastructure of this phenotype, we infected HEp-2 cells with strain JNEC-KY208 (O91:H–/Hg14, *stx1*^+^ *eibG*^+^) and examined the samples using scanning electron microscopy (SEM). SEM revealed that the bacteria remained firmly attached to the host cell surface, with pronounced constrictions at the interfaces between adjacent cells (Figure 1).

**Figure 1.**
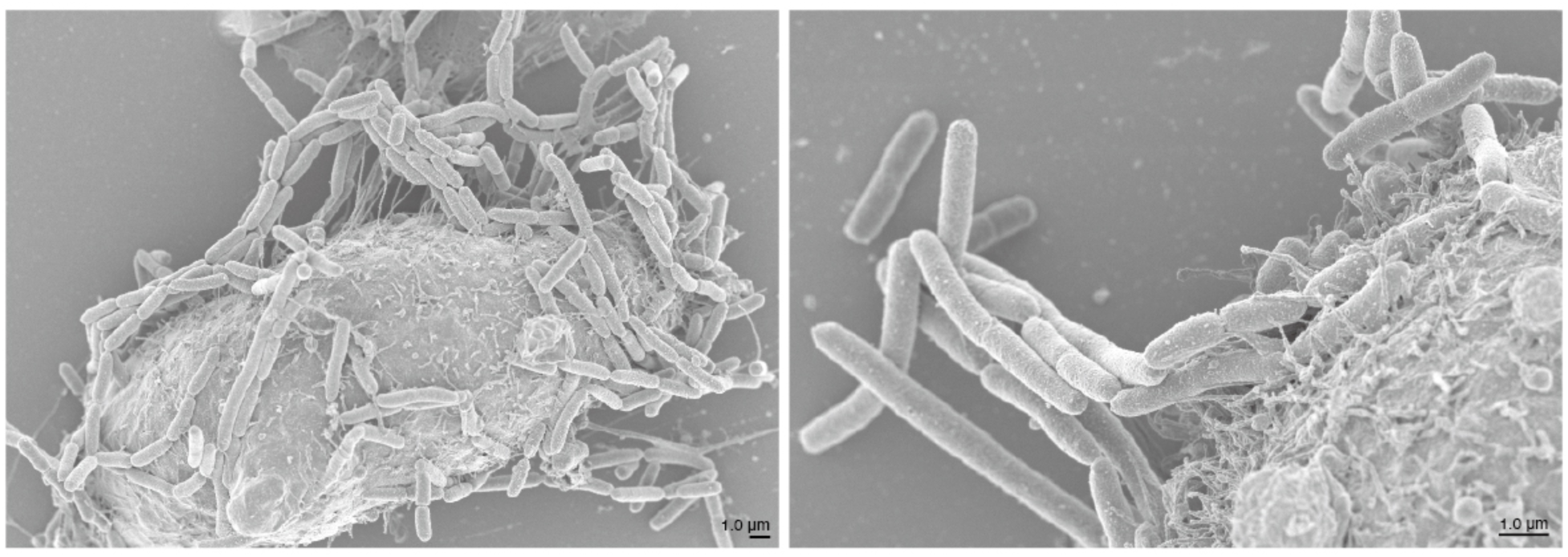
Scanning electron micrographs of HEp-2 cells infected with STEC strain JNEC-KY208. Cells were infected for 3 h under static conditions. Bacteria form multicellular chains that remain tightly attached to the host cell surface, displaying the characteristic chain-like adherence pattern (CLAP). Scale bars are indicated in each image.

Previous reports of CLAP have relied exclusively on static observations of infected HEp-2 cells at fixed time points and thus have not resolved how individual bacteria transition from single cells to multicellular chains. To visualize its dynamics, we performed time-lapse microscopy under static growth conditions. Approximately 60 min of dark-field imaging at 37°C showed that an individual bacterial cell elongated and subsequently underwent cell division without separation, giving rise directly to a chain formation (Figure 2A; Movie S1). Straightened image analysis demonstrated that constituent cells within a chain divided asynchronously, with an estimated doubling time of approximately ∼20 min (Figure 2B).

**Figure 2.**
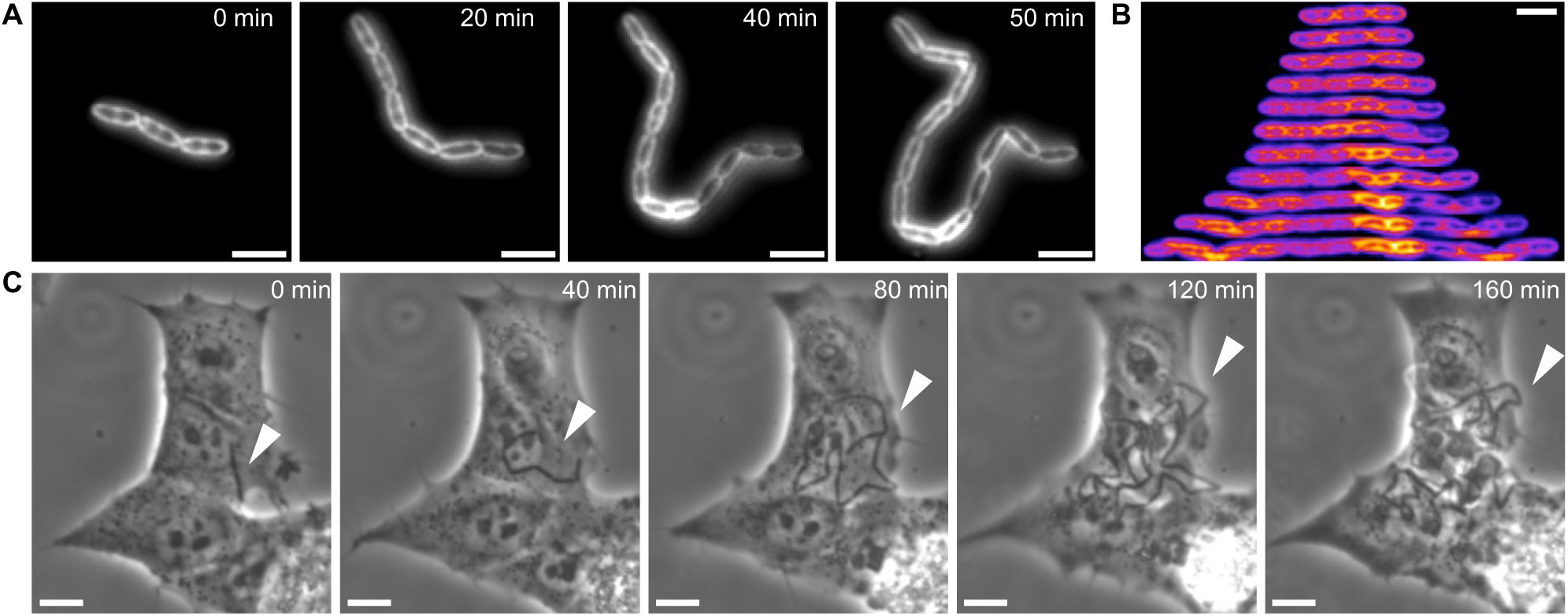
A single multicellular chain originates from a single cell in broth culture and during host-cell infection. (A) Time-lapse images of bacterial chain formation on a glass surface under dark-field microscopy at 37°C. Scale bar, 5 µm. (B) Straightened time-lapse images at 5 min intervals. The cells were straightened by image processing from panel A and are shown in pseudo color. Scale bar, 5 µm. (C) Time-lapse images of bacterial chain formation on HEp-2 cells under phase-contrast microscopy at 37°C. Scale bar, 10 µm. The white arrowhead indicates bacteria attached to the cell surface.

Next, we investigated whether chain formation occurred in the same manner during host infection, where CLAP was strongly manifested. To monitor infection dynamics, we established a 160 min label-free phase-contrast time-lapse imaging system at 37°C. HEp-2 cells on glass slides were infected with JNEC-KY208 at a multiplicity of infections (MOI) of 100, statically incubated for 1 h in 5% CO₂, and washed to remove non-adherent bacteria. Imaging revealed that adherent bacteria elongated and formed chains on host cells similarly to the observations in broth culture (Figure 2C; Movie S2).

### EibG-mediated multicellular chains function as a surface colonization strategy under flow conditions

However, the adaptive role of CLAP in physiological and environmental contexts remains unclear. Previous studies have described the characteristic bacterial dynamics under flow conditions that mimic natural and host-associated environments ^24,25^. To assess how chain morphology responds to shear stress, we established a flow-chamber system coupled to a syringe pump that continuously supplied fresh medium at defined flow rates (Figure 3A). Bacteria were introduced into the chamber and allowed to adhere to the glass surface during a 20 min incubation. When medium was supplied at 0.1 µL/s for 10 min, single cells were almost swept away, whereas multicellular chains remained on the surface (Figures 3BC, Movie S3). This flow rate corresponds to a shear stress of 0.06 Pa and a near-surface flow speed of 22 µm/s (Figure S1). Thus, chain morphology confers enhanced resistance to shear compared with individual cells.

**Figure 3.**
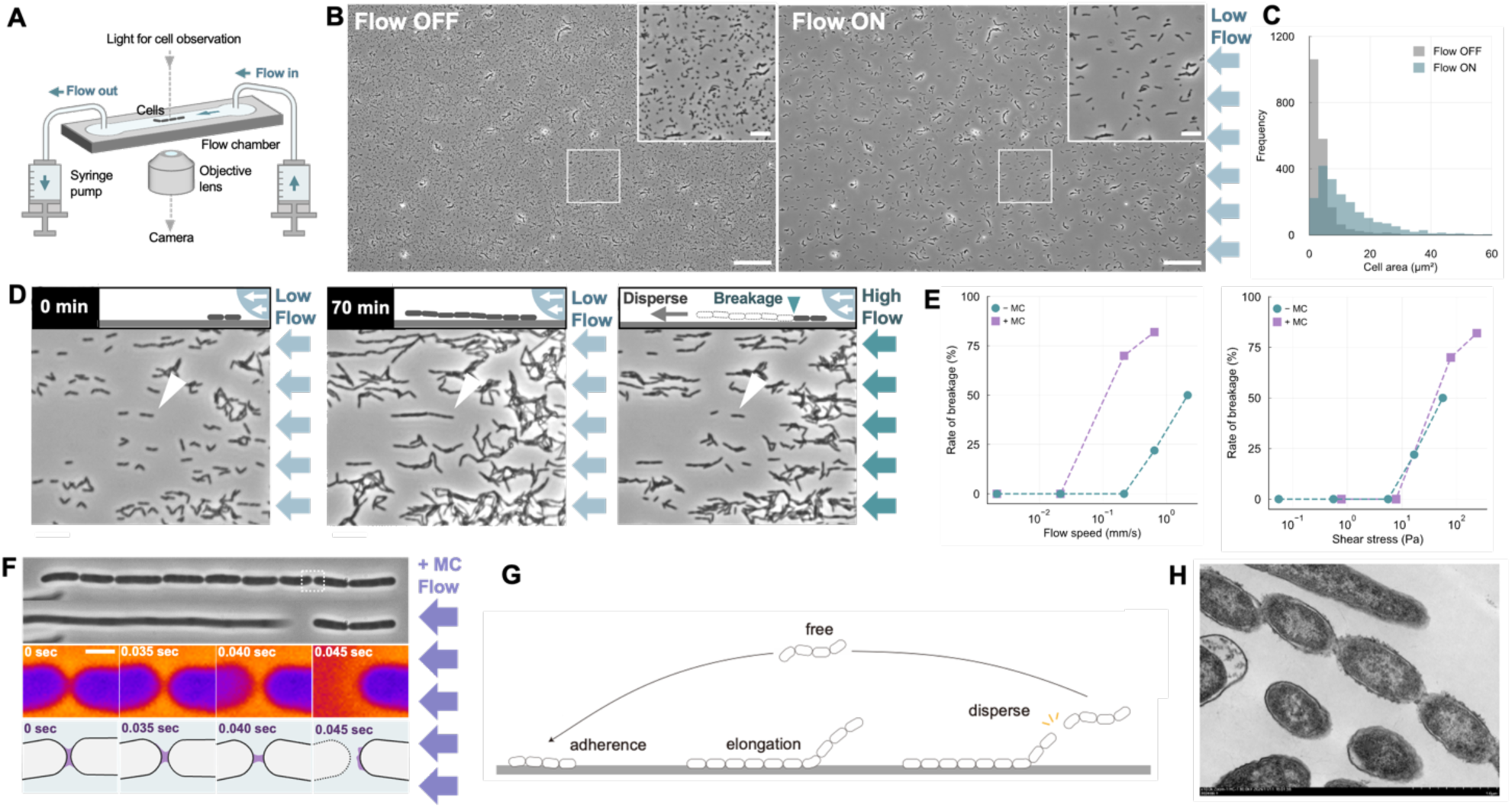
Multicellular chains function as a surface-colonization strategy under flow. (A) Schematic diagram of the experimental setup. The medium flow was applied from the right side of the fluid chamber using a syringe pump. Cell behavior was observed at room temperature. (B) Phase-contrast microscopy images before and after applying a water flow of 22 µm/s for 10 min. The white square indicates the region enlarged in the insets. Scale bars, 100 µm for the main images and 20 µm for the insets. (C) Distribution of cell area shown in panel B (N = 2,000 cells). (D) Chain formation and dispersion under water flow. Cells before applying a water flow (left), after applying a water flow of 22 µm/s for 70 min (center), and after applying a water flow of 22 mm/s for 1 min (right). The inset shows a schematic diagram of chain breakage and dispersion downstream. Scale bar, 20 µm. (E) Relationship between cell breakage and flow conditions. The ratio of broken cells among chain-forming cells was plotted against flow speed (left) and shear stress (right). Breakage during a 1-min flow of chain-forming clusters consisting of more than three connected cells was examined under conditions with or without 0.5% methylcellulose (MC) (N = 50 cells for each flow speed and shear stress). (F) High-speed imaging of chain breakage. Top: Phase-contrast microscopy images of chain breakage under a water flow of 2.2 mm/s with MC. Scale bar, 3 µm. The white square indicates the region enlarged in the middle panel. Middle: Enlarged time-lapse images of chain breakage shown in pseudo color. Scale bar, 0.5 µm. Bottom: Schematic images of chain breakage following the elongation of the intercellular distance. (G) Schematic model of the surface-colonization strategy inferred from microscopy. (H) Transmission electron micrographs of multicellular chains.

To examine how longer chains respond to higher shear forces, bacteria were first grown under low-flow conditions (0.1 µL/s) within the chamber to allow formation of extended chains. Upon applying a flow rate of 100 µL/s, chains ruptured at intercellular constriction sites and released small fragments downstream (Figure 3D; Movie S4). To test the dependence of cell breakage on shear stress, we observed cell breakage under different viscosity conditions, with and without 0.5% methylcellulose (MC) (Figure 3E, Movie S5). Even at the same flow speed, the presence of MC resulted in a higher rate of cell breakage, indicating that cell breakage depended on shear stress rather than shear rate. Notably, the force required to disrupt cell-cell junctions was lower than that required to detach the chains from the surface, indicating that fragmentation occurred prior to detachment (Movie S6). This provides an efficient means for disseminating clonal bacteria downstream.

High-speed imaging was performed at the moment of cell breakage (Figure 3F, Movie S7). The intercellular distance gradually increased and then suddenly ruptured at a certain point. These observations suggest that the breakage of chain-forming cells occurs due to the mechanical rupture of EibG-mediated connections at the poles of divided cells. Chain-mediated dispersal resembles recent observations in Enterococcus, where multicellular chains modulate colonization dynamics in response to flow^24^. Our findings demonstrate that gram-negative STEC can similarly exploit chain architecture: EibG-mediated adhesion promotes chain elongation, followed by shear-induced fragmentation, thereby expanding the spatial range of colonization (Figure 3G). Transmission electron microscopy confirmed that cytoplasmic separation between adjacent cells was complete, strongly suggesting that the fragmented units are viable and capable of establishing new foci (Figure 3H).

### EibG exhibits considerable sequence diversity

To investigate the conservation of the adhesins responsible for this phenotype, we chose 13 Japanese clinical STEC isolates as described in Materials and Methods. Together with JNEC-KY208, these isolates were subjected to whole-genome sequencing using the Illumina short-read and MinION long-read technologies. Hybrid assemblies generated using Hybracter yielded complete circular chromosomal contigs for all strains (Table 1). CheckM2 confirmed high-quality genomes, with completeness of 100% and contamination of ≤0.94%. In addition to Shiga toxin, various virulence factors were identified in these strains (Figure S2).

**Table 1.**
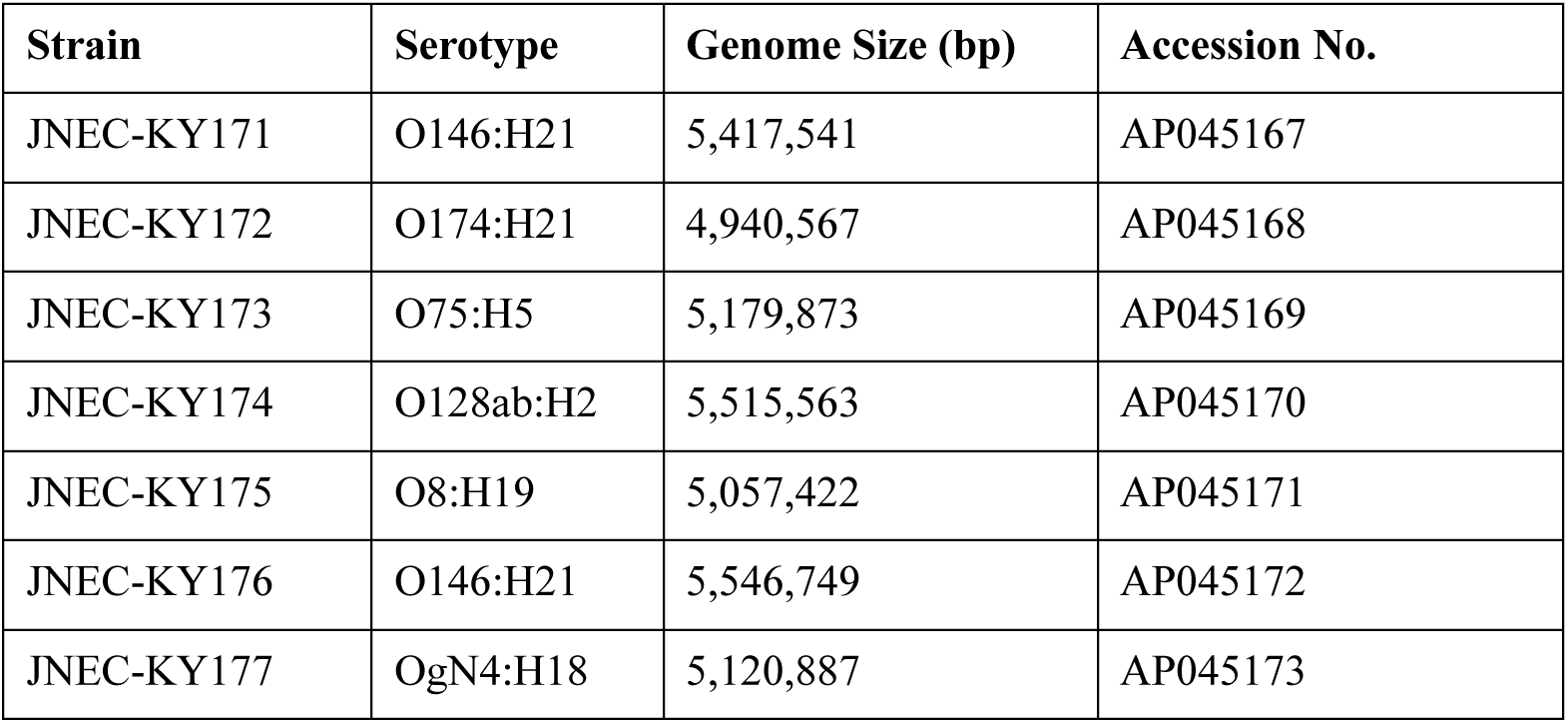
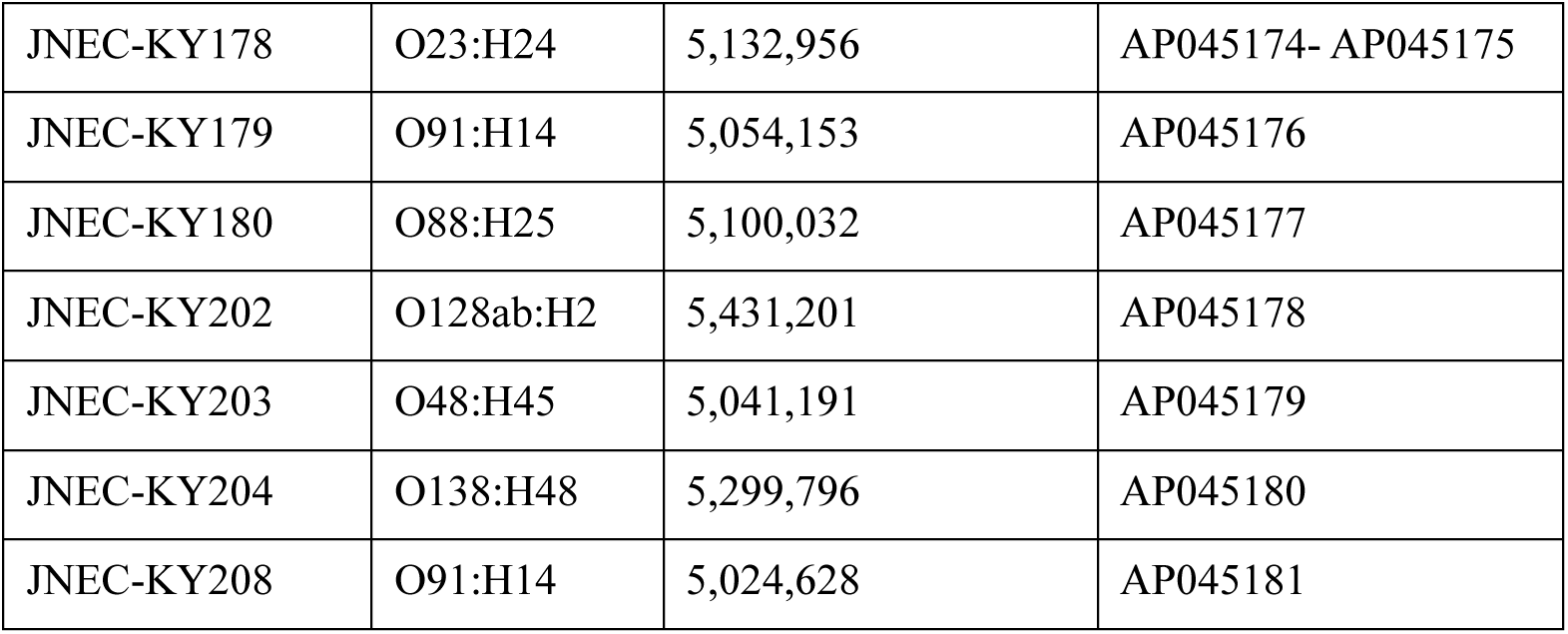
Complete genome sequences of 14 clinical STEC isolates determined in this study.

Using the 14 complete genomes as a database, we performed BLASTp searches with EibG (Accession ADJ17706.1) as the query. We identified 55 EibG-like amino acid sequences. After removal of signal peptides using SignalP, sequences were aligned with MAFFT, trimmed with trimAl, and subjected to phylogenetic inference using IQ-TREE3. Sequences lacking signal peptides were excluded, resulting in 41 EibG-like proteins used for further analysis. Interestingly, several sequences clustered distantly from known Eib-family proteins and instead grouped more closely with Saa, an autoagglutinating adhesin in STEC (Figure 4A). Because the functions of these proteins are unknown, we designated the two divergent clades as Yal (function unknown adhesin-like). Genomic context analysis was performed to explore potential evolutionary origins. Using clinker, we visualized loci spanning 10 kb upstream and downstream of each *eibG*-like gene. Known Eib-family proteins fell into three major genomic-context patterns, whereas the newly identified Cla (blue) and Yal (yellow) loci exhibited distinct, non-overlapping arrangements (Figure 4B). These results suggest that Cla and Yal are evolutionarily distinct from canonical Eib-family proteins and may represent separate adhesin lineages.

**Figure 4.**
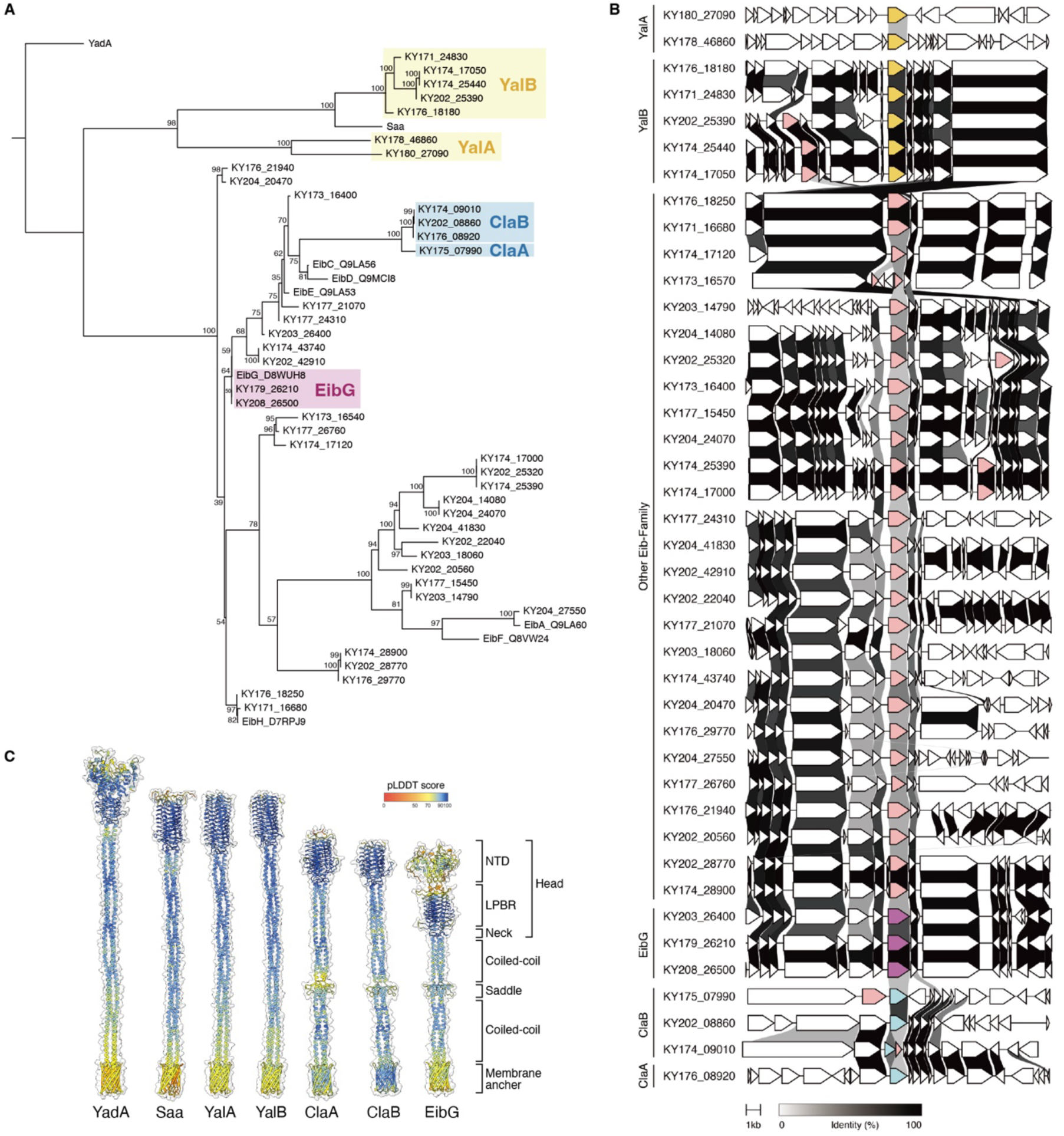
Phylogenetic, genomic, and structural analysis of EibG-like oligomeric coiled-coil adhesins. (A) Maximum-likelihood phylogeny of 41 EibG-like adhesins identified from 14 complete genomes of LEE-negative STEC and reference sequences. Signal peptides were removed by SignalP prior to alignment, and sequences were aligned using MAFFT and trimmed with trimAl. Tree inference was performed using IQ-TREE with ultrafast bootstrap support. Selected substitution model by ModelFinder is VT+F+G4. YadA was used as the outgroup to root the tree. The phylogeny includes two distinct adhesin groups identified in this study. These groups are indicated in the figure as Cla, shown in blue, and Yal, shown in yellow. (B) For each *eibG*-like gene, syntenic regions spanning 10 kb upstream and downstream are shown, visualized using clinker. The sequence homology is displayed in grayscale. (C) AlphaFold3 structural predictions of trimeric assemblies of representative adhesins. The sequence with the signal peptide removed by SignalP was used as input. The cartoon representation was color-coded according to the pLDDT score. NTD indicates the N-terminal domain, whereas LPBR indicates the left-handed parallel β-roll domain.

To compare structural similarities, we performed trimeric structural prediction using AlphaFold3, informed by the fact that OCA-family adhesins function as trimers. The newly identified Cla and Yal proteins displayed trimeric architectures that closely resembled the lollipop-like structures that are characteristic of YadA and EibG (Figure 4C). Cla proteins exhibited a pronounced saddle domain, consistent with structural predictions for EibG. In contrast, the saddle domain was not predicted in YadA, Saa, or the Yal proteins. Furthermore, the NTD predicted for EibG was absent in both Cla and Yal, indicating that although these proteins share an overall OCA-type trimeric structure, they differ in key structural features conserved among canonical Eib-family members.

### ClaA and ClaB are newly identified CLAP factors

To investigate the functional roles of ClaA and ClaB, we cloned each gene and expressed them in the JNEC-KY208 Δ*eibG* mutant. As expected, the Δ*eibG* strain exhibited a complete loss of CLAP, which was rescued by plasmid expression of EibG. Remarkably, expression of either ClaA or ClaB also restored CLAP (Figure 5A). However, when bacteria were grown under static culture conditions without host cells, ClaA-expressing cells formed few or no chains, whereas EibG-expressing cells formed multicellular chains and exhibited auto-aggregated (Figure 5B).

**Figure 5.**
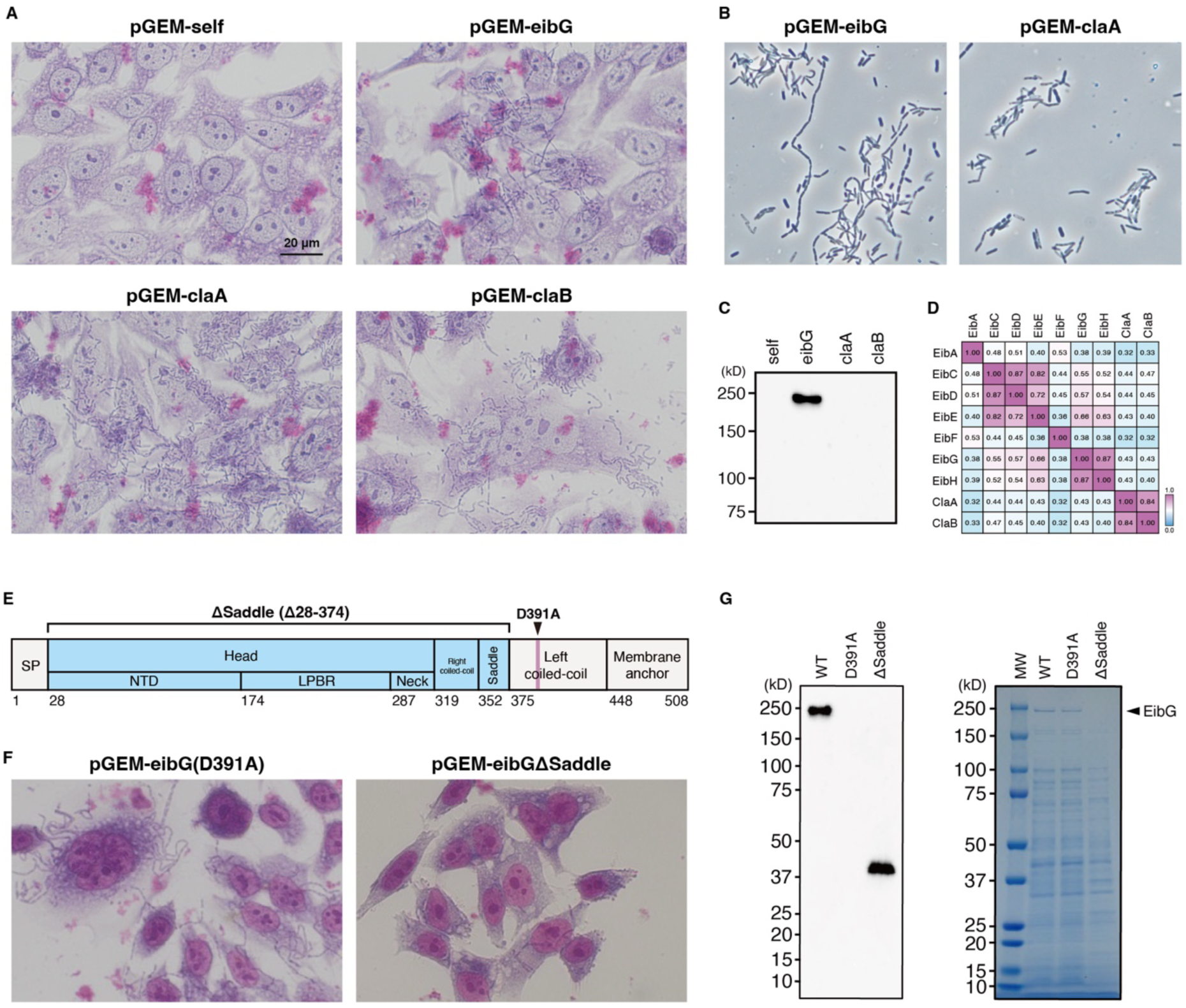
Functional characterization of Cla adhesins and structural separation of chain formation and IgG binding in EibG. (A) Complementation analysis of CLAP in the *eibG* deletion mutant by by plasmid-borne *eibG*, *claA*, and *claB*. Image shows that deletion of *eibG* abolishes CLAP, which is restored by plasmid-borne expression of *eibG*. The JNEC-KY208Δ*eibG* strain was infected with HEp-2 cells using a strain in which *eibG*, *claA*, and *claB* were complemented from a pGEM plasmid and *eibG* upstream region (168 bp). After a 3 h incubation, the cells were fixed and stained with Giemsa. Scale bar is 20 µm (B) JNEC-KY208 Δ*eibG* strain was cultured overnight under static conditions after introducing pGEM-*eibG* and pGEM-*claA*. (C) Immunoblotting assay of EibG, ClaA and ClaB. Whole-cell lysates were separated by SDS-PAGE and probed with HRP-conjugated human IgG Fc (20 ng/mL). High-molecular-weight band is detected only in strains expressing EibG. The label for each sample indicates the gene cloned into pGEM. (D) Sequence similarity matrix of Eib-family proteins and Cla adhesins. Pairwise amino acid sequence similarities were calculated using Clustal Omega and visualized as a heat map. Color scale indicates percent identity. (E) Domain architecture of EibG and positions of engineered mutations. The D391A substitution targets the residue aligned with the conserved IgG-binding aspartate in EibE. The Δsaddle mutant lacks the entire head–to–saddle region, which is indicated in the schematic by the solid line marking the deleted segment. SP indicates the Signal peptide, whereas NTD indicates the N-terminal domain and LPBR indicates the left-handed parallel β-roll domain. (F) Complementation analysis of CLAP in the *eibG* deletion mutant by by plasmid-borne *eibG* (D391A), *eibG* (ΔSaddle). HEp-2 infection assays were performed as in panel A. (G) Immunoblot and total-protein analysis of EibG mutants. Whole-cell lysates from strains expressing the indicated EibG variants were separated by SDS-PAGE and analyzed either by IgG Fc probing (left) or by total protein staining (right). The D391A mutant lacks detectable IgG Fc binding but retains the high-molecular-weight EibG band in the total-protein gel. In contrast, the Δsaddle mutant preserves IgG-binding activity, indicating that the saddle region is not required for Fc interaction. The labels for each sample indicate the eibG mutation cloned into pGEM, with WT representing wild-type EibG.

Since Eib proteins exhibit IgG Fc-binding activity, we examined whether Cla proteins also possess this activity. For this experiment, whole-cell lysates were prepared from unwashed HEp-2 infection cultures, allowing recovery of both HEp-2-associated and non-adherent bacteria together with the host cells. These lysates were boiled, separated by SDS-PAGE, transferred onto PVDF membranes, and probed directly with HRP-conjugated human IgG Fc fragment. In the strain transformed with pGEM-*eibG*, a prominent high molecular weight polypeptide corresponding to EibG was detected as an IgG Fc-binding band (Figure 5C). In contrast, no IgG-binding signal was detected in cells expressing ClaA or ClaB (Figure 5C). These results were further corroborated by the observation that the residues predicted to mediate IgG binding were not conserved in Cla (Figure S3). Collectively, these findings indicate that ClaA and ClaB lack IgG Fc-binding activity, in contrast to EibG.

To determine whether ClaA and ClaB should be considered members of the Eib-family proteins, we evaluated their sequence relatedness similarity matrix using Clustal Omega. Eib-family proteins formed several similarity clusters (EibA; EibC–E; EibF; and EibG–H), consistent with their established phylogenetic relationships (Figure 5D). In contrast, ClaA and ClaB did not cluster with any of these groups and shared only limited pairwise similarity with Eib proteins (>50%). Their distinct genomic contexts, lack of IgG Fc-binding activity, and limited similarity to canonical Eib proteins indicate that ClaA and ClaB form an adhesin lineage separate from the Eib family and are responsible for promoting CLAP. We therefore designate this lineage as chain-like adhesins (Cla).

To investigate whether *claA* and *claB* are present outside the Japanese clinical isolates analyzed in this study, we focused on a well-defined surveillance collection of LEE-negative STEC isolates from England^26^. In this cohort, serotypes O128:H2, O91:H14 and O146:H21 were among the top five STEC serotypes isolated from patients with gastrointestinal symptoms between 2014 and 2022. Using the accession numbers provided, we retrieved 1,354 draft genomes that were publicly available and screened them using BLASTn, applying thresholds of >90% nucleotide identity and >90% query coverage to detect *eibG*, *claA*, and *claB*. Strikingly, *claB* was present in 97.82%, 91.67%, and 98.26% of isolates belonging to serotypes O128:H2, O91:H14, and O146:H21, respectively (Table 2).

**Table 2.** Frequency of *claA*, *claB* and *eibG*.

|  | *claA* | *claB* | *eibG* |
| --- | --- | --- | --- |
| O128:H2 | 0/275<br>(0.00%) | 269/275<br>(97.82%) | 2/275<br>(0.73%) |
| O91:H14 | 0/492<br>(0.00%) | 451/492<br>(91.67%) | 7/492<br>(1.42%) |
| O146:H21 | 0/518<br>(0.00%) | 509/518<br>(98.26%) | 0/518<br>(0.00%) |
| Other | 0/69<br>(0.00%) | 65/69<br>(94.20%) | 0/69<br>(0.00%) |
| Total | 0/1354<br>(0.00%) | 1294/1354<br>(95.57%) | 9/1354<br>(0.66%) |

### Distinct structural domains mediate CLAP and IgG binding in EibG

Next, we examined the domain architecture underlying the dual functions of EibG: CLAP formation and IgG Fc-binding activity. Previous structural studies of EibE and multiple sequence alignments of Eib-family proteins have identified two residues (EibE^D378^–^Y388^) that are critical for interaction with the IgG Fc region, although their roles in CLAP have not been defined ^20^. In addition, deletion of the saddle domain disrupts chain formation; however, whether this region contributes to IgG Fc-binding has remained unclear ^23^.

To experimentally test the contribution of the predicted IgG Fc-binding site to CLAP, we mapped the residues identified in EibE ^D378^–^Y388^ onto the EibG sequence. In EibG, the position corresponding to EibE^Y388^ is already an alanine (EibG^A392^), whereas EibE^D378^ aligns with EibG^D391^. We therefore generated an EibG^D391A^ mutant to disrupt the conserved aspartate predicted to participate in IgG Fc binding activity (Figure 5E). In parallel, we constructed a Δsaddle mutant lacking the entire saddle domain (Figure 5E). The EibG^D391A^ point mutant retained CLAP formation, and a high molecular weight EibG band was still detectable by SDS–PAGE, but no IgG Fc-binding signal was observed (Figures 5F, 5G). In contrast to the EibG^D391A^ point mutant, the Δsaddle mutant showed loss of CLAP while retaining IgG Fc-binding activity (Figures 5F, 5G). These results demonstrate that the head-to-saddle domain and IgG Fc-binding residues contribute independently to EibG function, establishing that chain formation and IgG binding are mediated by distinct, non-overlapping structural determinants.

### EibG functions as a virulence factor in vivo

EibG is frequently detected in STEC and contributes to adhesion; however, its role in pathogenicity has not been fully defined. To determine whether EibG contributes to virulence in vivo, we used an mouse intraperitoneal (i.p.) infection model using specific pathogen-free (SPF) mice. We adopt this model because it provides a clear and reproducible means of evaluating the systemic impact of STEC. Lethality provides a clear and quantifiable endpoint, allowing the evaluation of pathogenic consequences without the confounding variability inherent in gastrointestinal colonization models^27,28^.

Because Shiga toxin is the principal virulence determinant in STEC, we constructed a JNEC-KY208Δ*eibG* to allow assessment of EibG-specific effects. Mice were randomly assigned to receive 1.5 × 10⁸ CFU of either the Δ*stx1* parental strain or the isogenic Δ*stx1*Δ*eibG* mutant by intraperitoneal injection and were monitored for five days. Across three independent experiments (n = 31 for Δ*stx1*; n = 25 for Δ*stx1*Δ*eibG*), the Δ*eibG* mutant exhibited a significantly reduced lethality compared with the parental strain (*p*=0.044) (Figure 6). These findings demonstrate that EibG makes a clear contributes to STEC virulence in vivo.

**Figure 6.**
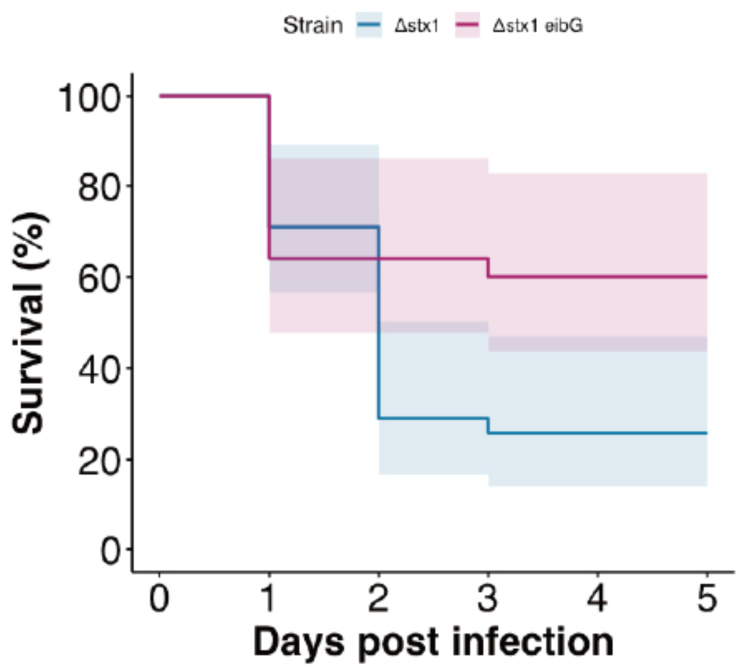
EibG contributes to virulence in a mouse infection model. Survival curves of mice infected intraperitoneally with JNEC-KY208 derivatives. Specific-pathogen-free mice were inoculated intraperitoneally with 1.5 × 10⁸ CFU of either the Δ*stx1* strain or the isogenic Δ*stx1*Δ*eibG* strain. Animals were monitored for 5 days for survival outcomes. Across three independent experiments (n = 31 for Δ*stx1*; n = 25 for Δ*stx1*Δ*eibG*), deletion of *eibG* resulted in significantly reduced lethality compared with the parental strain (log-rank test, *p* = 0.044).

## Discussion

In this study, we elucidate the genetic, structural, and functional basis of the CLAP observed in certain LEE-negative STEC. By integrating live-cell imaging, comparative genomics, and targeted mutagenesis, we demonstrated that EibG-driven multicellular chains constitute a previously unrecognized surface colonization strategy for STEC. Our findings indicate that chain morphology enhances adhesive persistence under dynamic conditions, promotes clonal dispersal through shear-induced fragmentation, and contributes to pathogenic potential in vivo. Furthermore, the discovery of a new class of adhesins (Cla) that mediates CLAP but lacks immunoglobulin-binding activity substantially broadens our current understanding of host-surface attachment mechanisms across DEC.

A central advancement of this study is the temporal resolution of CLAP development. Previous reports relied on static snapshots of infected cells and did not resolve whether CLAP arises through filamentous elongation, aggregation, or modified cell division. Our time-lapse analyses clearly show that chains originate from single progenitor cells that elongate and divide without separation. By integrating this imaging approach with microfluidic flow assays, we found that chain morphology provides an adaptive advantage under shear conditions. Solitary cells detached readily under flow, whereas multicellular chains remained adherent and fragmented at intercellular constrictions, thereby releasing viable progeny downstream. This dispersion-driven mode of colonization parallels recent observations in *Enterococcus faecalis*, in which multicellular chains promote flow-dependent niche expansion^24^. Our results extend this paradigm to Gram-negative pathogens, suggesting that chain-based architectures represent a convergent strategy among host-associated bacteria that must maintain attachment and spread within fluid-exposed environments.

For DEC inhabiting the gastrointestinal tract, mucus flow and peristalsis generate substantial shear that can disrupt surface attachment^29,30^. Although higher shear forces promote more efficient chain fragmentation, our data indicate that chain breakage occurs at shear stresses of 0.78 Pa for observations more than 1 h (Movie S4). This value is broadly comparable to previously estimated shear stresses in the intestinal lumen and is therefore consistent with the possibility that such fragmentation may also occur in vivo^31^. Our results highlight a potential infection strategy that may be exploited by certain LEE-negative STEC in the intestinal environment. In this context, chain formation provides a significant advantage in terms of fitness. Accordingly, our findings add flow-dependent colonization to the repertoire of biological activities previously attributed to OCAs, which including adherence, biofilm formation, invasion, intracellular survival, serum resistance, and cytotoxicity.

Comparative genomic analysis revealed unexpected diversity within EibG-like adhesins. Although EibG and its homologues have historically been classified within the immunoglobulin-binding protein family, our phylogenetic analyses identified distinct clusters with unique genomic contexts, forming two previously unrecognized lineages that we designate Cla and Yal. This diversification aligns with the known evolutionary plasticity of OCA-family proteins, which frequently undergo domain shuffling, repetition, and recombination, and are mobilized through horizontal gene transfer processes^32^. Structural predictions provide further evidence for the evolutionary divergence of Cla proteins from canonical Eib-family members. Despite adopting trimeric OCAs, Cla proteins do not exhibit key features typically associated with the Eib family, such as conserved immunoglobulin G (IgG) Fc-binding residues and an NTD. This structural deviation suggests a functional specialization of Cla proteins in host-cell adherence rather than an interaction with immunoglobulins.

This functional divergence was corroborated by genetic analyses. Expression of ClaA or ClaB restored CLAP in the *eibG* deletion mutant, demonstrating that Cla proteins are sufficient to functionally complement the loss of EibG for this phenotype. However, under static culture conditions, Cla-expressing strains showed minimal chain formation, implying that Cla-dependent CLAP may be specifically displayed under host-associated or otherwise specialized environmental conditions. Notably, Cla proteins lack IgG Fc-binding activity, which directly establishes that CLAP and immunoglobulin binding are mechanistically separable phenotypes driven by distinct subsets of the OCA family proteins. A detailed examination of the domain architecture of EibG provided additional resolution for this functional separation. Substitution of the predicted IgG binding residue EibG^D391A^ abolished Fc recognition while leaving CLAP intact, whereas removal of the saddle domain eliminated CLAP while preserving IgG-binding. These results demonstrate that these two activities are encoded by distinct structural modules within EibG. These results highlight the modular organization of OCA adhesins and raise broader questions about the selective forces that preserve the immunoglobulin-binding capability within the Eib lineage.

Consistent with this hypothesis, ClaB was highly prevalent among clinical LEE-negative STEC isolates from England, particularly serotypes O128:H2, O91:H14, and O146:H21, which were among the most frequently reported during 2014-2022. The high prevalence of ClaB in these clinically important lineages underscores its importance as an adhesin and suggests that Cla-mediated chain-like adhesion may contribute to its prevalence among clinical isolates. Our findings suggest that the chain-formation function of OCA proteins may provide a fitness advantage under specific ecological conditions.

Our in vivo experiments revealed that EibG contributes to pathogenicity. Although Shiga toxin remains the major driver of STEC-associated mortality, the *eibG* deletion mutant exhibited significantly reduced lethality even in an Δ*stx1* background. This indicates that EibG enhances disease severity through a toxin-independent mechanism. Potential explanations include improved colonization efficiency, increased persistence at host surfaces, or modulation of innate immune responses. While our current data do not resolve whether CLAP, IgG binding, or both phenotypes underlie the virulence effect, the domain analyses suggest that the physical properties of chain formation could themselves influence infection dynamics. Future studies employing domain-specific EibG mutants will be essential for dissecting host responses to these modular architectural features.

Collectively, our study expands the conceptual framework governing adhesion and virulence in STEC. Chain-like adherence, enabled by genetically diverse OCA family adhesins with distinct structural and functional properties, has emerged as a key strategy for surface colonization in certain LEE-negative STEC strains. The identification of Cla proteins underscores the evolutionary capacity of DEC lineages to acquire and repurpose adhesion modules via horizontal gene transfer. A deeper understanding of the distribution, regulation, and host interactions of EibG, Cla, and related adhesins is crucial to predict predicting the pathogenic potential of emerging LEE-negative STEC and hybrid DEC lineages. Moreover, these findings highlight OCA-mediated attachment as a promising target for therapeutic and preventive intervention.

## Materials and Methods

### Bacterial strains, media, primers and plasmids

All resources used in this study are listed in Tables S1 (strains), S2 (primers), and S3 (plasmids). Bacteria were grown at 37℃ in LB Lenox medium (BD, USA) supplemented with ampicillin (100 μg/mL), chloramphenicol (20 μg/mL), kanamycin (50 μg/mL).

### Genetic Manipulation of STEC strains

Genetic manipulation of STEC strains was performed using the λ Red recombination method with the pSIM6 plasmid^33^. The *eibG* gene, including 168 bp of its upstream region, was cloned into pGEM-T Easy (Promega, USA). The pGEM-self was used as a negative control^34^. For *claA* and *claB* constructs, the *eibG* coding sequence (CDS) was replaced with each CDS via seamless cloning, preserving the *eibG* upstream region. The X-cocktail method, which relies on Exonuclease III (Takara Bio, Japan), was used as previously described^35^. Inverse PCR for site-directed mutagenesis was performed using partially overlapping primers and KOD One DNA polymerase (TOYOBO, Japan).

### Cell culture and infection with bacteria

HEp-2 cells were grown in Minimum Essential Medium (Sigma-Aldrich, USA) supplemented with FBS (10%) (Thermo Fisher Scientific, USA) at 37°C in 5% CO_2_. The procedures for observing STEC adherence have been described previously ^17^.

### SEM

The samples were fixed overnight at 4 °C with a mixture of 2.5% glutaraldehyde, 2% paraformaldehyde, and 0.5% acetic acid. After fixation, the samples were dehydrated using a graded acetone series (50–99.5%) at room temperature and dried using a critical point dryer with CO₂ (CPD300, Leica Microsystems, Germany). The dried samples were coated with osmium using an osmium plasma coater (Neoc-Pro/P, Meiwafosis, Japan) and subsequently examined with a field-emission SEM (FE-SEM; SU8600, Hitachi High Technologies, Japan).

### Optical microscopy and data analyses

For time-lapse imaging of chain formation, bacteria and HEp-2 cells on the glass surface were visualized under an inverted microscope (IX83; Olympus, Japan) equipped with an ×40 objective lens (LUCPLFLN 40×PH, N.A. 0.6), a CMOS camera (DMK33UX174; Imaging Source, Germany), and an optical table (ASD-1510T). Samples were visualized using a halogen lamp (U-LH100L-3; Olympus, Japan) through a bright-field condenser (IX2-LWUCD, N.A. 0.55; Olympus, Japan) for phase-contrast microscopy, or a dark-field condenser (U-DCD, NA0.8-0.92; Olympus, Japan) for dark-field microscopy. The microscope stage was heated using thermoplate (TP-110R-100; Tokai Hit, Japan). Projections of the images were acquired with the imaging software IC Capture (Imaging Source) under at 10-s resolution and converted into an AVI file without any compression. All data were analyzed by Fiji software (ImageJ 1.53t).

For the flow experiment, the samples were examined under the inverted microscope equipped with ×10 and ×100 objective lenses (UPLFLN 10×2PH; N.A. 0.3, UPLXAPO100×OPH, N.A. 1.45; Olympus, Japan), a CMOS camera, and an optical table. The projection of the image was captured with the imaging software IC Capture at the resolutions ranging from 0.005-s to 1-s and converted into an AVI file without any compression.

### Time-lapse imaging of chain formation

For the observation of bacterial chain formation on the glass surface, a tunnel chamber was assembled by taping coverslips with double-sided tape (∼10 µm thick; 7070W, Teraoka, Japan), slightly modified from a previous report^25^. Bacterial suspensions in LB medium were introduced into the chamber and incubated at 37°C for 10 min on a stage equipped with a thermoplate. Unattached cells were washed away with LB medium and the ends of the chamber were sealed with Vaseline.

For the observation of bacterial chain formation on HEp-2 cells, HEp-2 cells were seeded onto a coverslip in a Petri dish and incubated overnight at 37°C in 5% CO₂. The coverslip with HEp-2 cells was washed with FBS-free medium and incubated for 1 h at 37°C in 5% CO₂. Subsequently, a bacterial suspension in LB medium was added and incubated for 1 h at 37°C in 5% CO₂. The coverslip with HEp-2 cells was then washed twice with FBS-free medium and assembled into a tunnel chamber using double-sided tape (∼90 µm thick; NW-5, Nichiban, Japan). The ends of the chamber were sealed with Vaseline.

### Flow experiment

The flow chamber was assembled by taping a coverslip to a glass slide, as described previously^25^. The glass slide was bored using a high-speed drill press equipped with a diamond-tipped bit (1 mm diameter, No. 13853; NAKANISHI, Japan) for the inlet and outlet ports. The central portion of the double-sided tape (7082; Teraoka, Japan) was cut to a length of 25 mm long and 1.5 mm wide. The chamber was assembled by taping a coverslip onto a glass slide. The inlet and outlet ports (N-333; IDEX Health & Science, USA) were attached to a high-temperature elastic adhesive (Super X No. 8008 Clear, CEMEDINE, Japan). The finished channel of the sample chamber was straight with a width: 1.5 mm, height: 0.1 mm, and length: 25 mm for bright-field microscopy. A syringe pump (Legato 200; Kd Scientific, USA) was connected to the flow chamber by a connector and tube (F-333NX and 1512L; IDEX Health & Science, USA).

Bacterial suspensions in the LB medium were injected into the chamber and incubated for 15 min. The unattached cells were slowly washed away by LB medium with or without 0.5% methylcellulose (M0512, viscosity: 4,000 cP at 2%, Sigma-Aldrich, USA), at a flow rate of 0.1 µL/s for 1 h. The behavior of attached cells under fluid flow was observed at room temperature (RT). The flow rate of each syringe pump was calibrated to the velocity of the fluid flow near the glass surface at RT. The near-surface flow speed was determined from dispersed cells that moved passively in the flow direction following occasional breakage of multicellular chains at the glass surface. The shear stress at the chamber surface was estimated using the equation described previously ^36^.

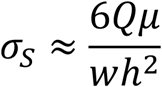

Where *σ_S_* is the shear stress at the chamber surface; *Q* is the flow rate; *µ* is the viscosity of the solution, assumed to be 1 mPa·s for LB medium without MC and 14 mPa·s for LB medium containing 0.5% MC; and *w* and *h* are the width and height of the flow chamber, respectively.

### Transmission electron microscope

Bacterial cells were washed three times with PBS and fixed with 2% paraformaldehyde and 2.5% glutaraldehyde in 0.1M phosphate-buffer (pH 7.4), for 24hrs at 4°C. The samples were then post-fixed in osmium fixation solution [1% (w/v) osmium tetroxide, 1.25% (w/v) potassium ferrocyanide, and 5 mM calcium chloride and embedded in 2% (w/v) low-melting-point agarose. After staining the blocks with 0.5% aqueous uranyl acetate, the specimens were dehydrated and embedded in Spurr resin. Thin sections were mounted on copper grids and post-stained with saturated uranyl acetate and lead citrate. The specimens were observed using an HT7700 transmission electron microscope (Hitachi High Technologies, Japan).

### Complete genome sequencing

Genomic DNA was extracted using the KingFisher Duo Prime System (Thermo Fisher Scientific, USA). For short-read sequencing, the extracted DNA was prepared library using the QIAseq FX DNA Library Kit (Qiagen, Germany) and paired-end (150 × 2) sequencing was performed on the Illumina platform. For long-read sequencing, libraries prepared using a Rapid Barcoding Kit V14 (Oxford Nanopore Technologies, UK) were sequenced on the MiniON Mk1B device. Sequencing reads were assembled using Hybracter v0.11.0^37^. Among the plasmid contigs obtained after assembly, only the plasmid containing the *eibG*-like gene was selected for inclusion in the final complete genome sequences. The quality of the assembled genomes was assessed using CheckM2 v1.1.0 ^38^. Gene annotation was performed using the DFAST platform provided by DNA Data Bank of Japan (https://dfast.ddbj.nig.ac.jp)^39^.

### Phylogenetic analysis of EibG-like proteins

EibG-like proteins were searched in the in-house clinical isolates genome database (not publicly available) using BLAST+ v2.16.0^40^. Based on these results, 14 representative strains harboring EibG-like protein candidates were selected and subjected to complete genome sequencing. EibG-like proteins were re-identified by BLASTp searches using the complete genome sequences of the 14 strains in the database. After removing signal peptides using SignalP v6.0^41^ (https://services.healthtech.dtu.dk/services/SignalP-6.0/), the identified EibG-like proteins, Saa, and YadA were aligned using MAFFT v7.525^42^, processed with trimAl v1.5^43^, and subsequently used to construct a phylogenetic tree with IQ-TREE v3.0.1^44^.

### Visualization of genomic context

To visualization of genomic context, the target gene sequences were obtained, including those 10 kbp upstream and downstream. Gene cluster comparison figure was generated using clinker and clustermap.js v0.0.31^45^.

### In silico structure prediction

The three-dimensional structure of the EibG-like proteins was predicted using AlphaFold3 (DeepMind) via the AlphaFold Server (https://alphafoldserver.com)^46^. The amino acid sequence without the signal peptide removed by SignalP 6.0 was used as the input. The predicted structures were visualized using UCSF ChimeraX v1.10.1 ^47^.

### Immunoblotting

The culture medium was mixed with Sample Buffer Solution with 3-Mercapto-1,2-propanediol (×4) (Fujifilm, Japan) and heated at 98°C for 10 minutes. The protein solution was electrophoresed at 20 mA for 75 minutes on a SuperSep Ace, 5-20% gel (Fujifilm, Japan), then transferred to a PVDF membrane at 200 mA for 60 minutes. The membrane was blocked with EveryBlot Blocking Buffer (Bio-Rad, USA) for 5 minutes, then reacted with 20 ng/mL Peroxidase ChromPure Human IgG, Fc fragment (Jackson ImmunoResearch, USA).

### Animal experiment

Three-week-old female specific pathogen-free (SPF) BALB/c mice were purchased (CLEA Japan, Japan). A total of 56 mice were used for the infection studies. The animals were randomly divided into two experimental groups (n = 31 for O91 Δ*stx1* and n = 25 for O91Δ*stx1*Δ*eibG*) and housed in cages (5-6 animals per cage). They were provided with sterilized food, bedding, and water ad libitum throughout the experimental period.

Mice were challenged by intraperitoneal (i.p.) injection with a 150 μL bacterial suspension containing approximately 1.5 × 10^8^ colony-forming-units of either O91 Δ*stx1* or O91Δ*stx1*Δ*eibG*. Survival was monitored and recorded every 24 h for 5 days. The infection experiments were replicated at least three times independently.

## Author Contribution

Conceptualization: K.Y., I.S.; Data curation: K.Y.; Formal analysis: K.Y., U.N.; Supervision: N.D., A.Y., I.S.; Funding: K.Y., I.S.; Investigation: K.Y., U.N., I.T., L.K., I.N., K.H., K.M.; Methodology: K.Y.; Visualization: K.Y., U.N.; Writing – original draft; K.Y., U.N. I.T., K.H., K.M.; Writing – review & editing: K.Y., I.S.

## Supporting information

Supplemental Informations

## Acknowledgements

We are deeply grateful to the EHEC Working Group, Japan for isolating the bacterial strains. We thank Saomi Ozawa and Toshiaki Yamagishi at the NIID for technical assistance. We also thank Koichiro Tamura at the TMU for helpful discussions.

## Ethics declarations

### Animal experiments

The animal experiments were carried out according to a protocol approved by the Institutional Animal Care and Use Committee of The Jikei University.

### Data availability

The sequencing data presented in this study have been deposited in the DNA Data Bank of Japan, accession number PRJDB37405.

