## Supplemental Informations for "Oligomeric coiled-coil adhesins that drive chain-like adhesion diversify surface colonization strategies in Shiga toxin-producing *Escherichia coli*"

**Supplementary Figure S1.**
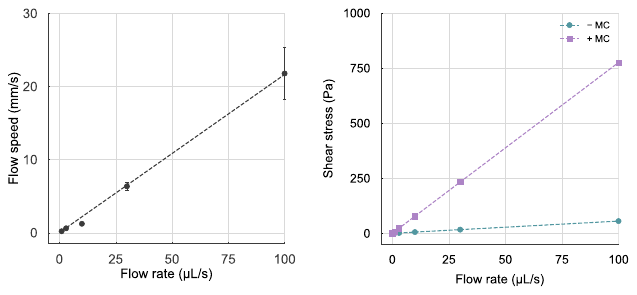
 **Calibration of flow dynamics**

Left: Relationship between the flow rate of the syringe pump and the flow speed near the glass surface, measured from the initial velocity of broken-chain cells detached from the surface (n = 3 cells at each flow rate). Right: Relationship between the flow rate of the syringe pump and shear stress under conditions with or without MC. Shear stress was calculated based on the geometry of the flow chamber, using viscosities of 1 mPa·s and 14 mPa·s for without and with MC, respectively.

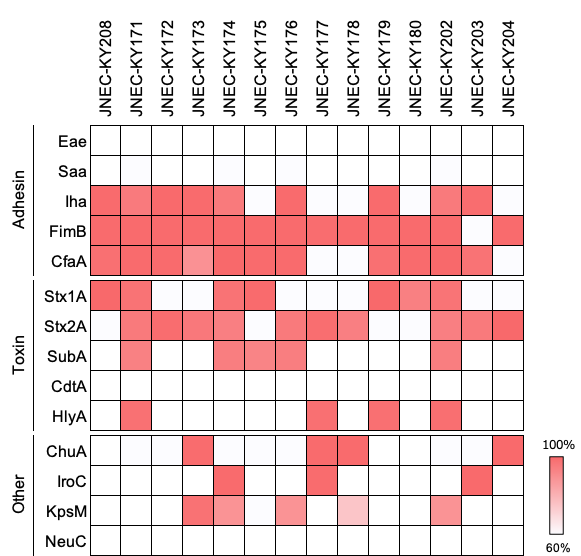

**Supplementary Figure S2. Virulence gene profiles of the 14 STEC isolates**

A panel of virulence-associated genes, including shiga toxin and additional DEC-related factors, was screened across all isolates. Gene presence and sequence similarity were assessed by BLAST, and percent identify values were visualized as a heatmap.

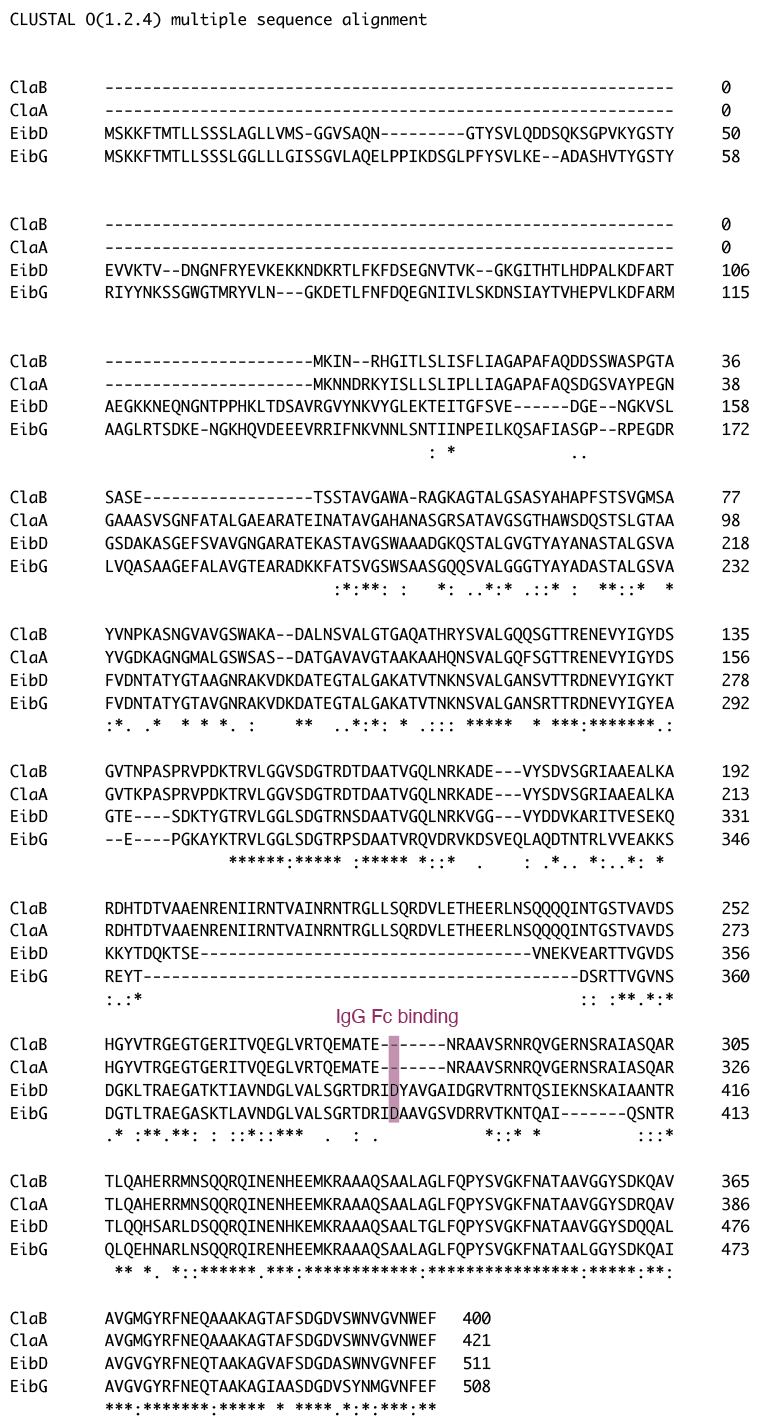

**Supplementary Figure S3. Multiple sequence alignment of the newly adhesins Cla and Eib**

The sequences of ClaA, ClaB, EibD, and EibG were aligned using CLUSTAL O (1.2.4). Residues involved in binding to IgG Fc are shown in pink.
