## Supplementary material for "Oligomeric coiled-coil adhesins that drive chain-like adhesion diversify surface colonization strategies in Shiga toxin-producing *Escherichia coli*": Movie Captions

**Movie Caption**

Movie S1. Time lapse imaging of chain formation. Bacterial cells on the glass surface were visualized under a dark-field microscopy at 37℃ for 1h.

Movie S2. Time lapse imaging of CLAP. Bacterial cells on HEp-2 cells surface were visualized under a phase contrast microscopy at 37℃ for 2.5 h.

Movie S3. Bacterial response to low flow under a low-viscosity condition. Bacterial cells were visualized by phase-contrast microscopy at room temperature (RT) for 1 h. Cells were injected into the flow chamber and incubated for 15 min to allow initial adhesion. The medium flow without MC was applied from the right side at the beginning of the movie. The flow rate was 0.1 µL/s, corresponding to a shear stress of 0.06 Pa at the surface and a near-surface flow speed of 22 µm/s.

Movie S4. Bacterial response to high flow under a low-viscosity condition. Bacterial cells were visualized by phase-contrast microscopy at RT. First, bacteria were grown under low flow conditions (22 µm/s) within the chamber to allow formation of extended chains. After 70 min incubation, high flows were applied at near-surface flow speeds of 6.5 mm/s and 22 mm/s.

Movie S5. Bacterial response to low flow under a high-viscosity condition. Bacterial cells were visualized by phase-contrast microscopy at RT for 1.5 h. Non-adherent cells were already washed away before imaging. The medium flow containing 0.5% MC was applied from the right side. The flow rate was 0.1 µL/s, corresponding to a shear stress of 0.78 Pa at the surface and a near-surface flow speed of 22 µm/s.

Movie S6. Chain breakage at high flow under a low-viscosity condition. Bacterial cells were visualized by phase-contrast microscopy at RT for 2 min. The medium flow without MC was applied from the right side at the time of 50 sec. The flow rate was 100 µL/s, corresponding to a shear stress of 55 Pa at the surface and a near-surface flow speed of 22 mm/s.

Movie S7. High-speed imaging of chain breakage at medium flow under a high-viscosity condition. Bacterial cell chain breakage was captured at 200 fps by phase-contrast microscopy at room temperature RT for 0.57 s. The medium flow containing 0.5% MC was applied from the right side. The flow rate was 10 µL/s, corresponding to a shear stress of 78 Pa at the surface and a near-surface flow speed of 2.2 mm/s. The purple arrowhead indicates the site of chain breakage. The latter half of the movie shows an enlarged view of the chain breakage event.
